# Regulatory cross-talk supports resistance to Zn intoxication in *Streptococcus*

**DOI:** 10.1101/2021.07.26.453799

**Authors:** Matthew J. Sullivan, Kelvin G. K. Goh, Glen C. Ulett

**Affiliations:** School of Medical Sciences, and Menzies Health Institute Queensland, Griffith University, Parklands, Australia 4222

**Author notes:** Correspondence: Professor Glen C. Ulett.

**Keywords:** metal ions, *Streptococcus*, copper, zinc, bacterial pathogenesis

## Abstract

Metals such as copper (Cu) and zinc (Zn) are important trace elements that can effect bacterial cell physiology but can also intoxicate bacteria at high concentrations. Discrete genetic systems for management of Cu and Zn efflux have been described in several bacteria pathogens, including streptococci. However, insight into molecular cross-talk between systems for Cu and Zn management in bacteria that drive metal detoxification, is limited. Here, we describe a biologically consequential cross-system effect of metal management in group B *Streptococcus* (GBS) governed by the Cu-responsive *copY* regulator in response to Zn. RNAseq analysis of wild-type (WT) and *copY*-deficient GBS exposed to metal stress revealed unique transcriptional links between the systems for Cu and Zn detoxification. We show that the Cu-sensing functions of CopY extend beyond Cu, and enable CopY to regulate Cu and Zn stress responses to effect genes involved in central cellular processes, including riboflavin synthesis. CopY also contributed to supporting GBS virulence *in vivo* following infection of mice. Identification of the Zn resistome of GBS using TraDIS revealed a suite of genes essential for GBS growth in metal stress. Several of the genes identified are novel to systems that support bacterial survival in metal stress, and represent a diversity of mechanisms of microbial metal homeostasis during cell stress. Overall, this study reveals a new and important mechanism of cross-system complexity driven by CopY in bacteria to regulate cell management of metal stress and survival.

**Author Summary:** Metals, such as Cu and Zn, can be used by the mammalian immune system to target bacterial pathogens, and consequently, bacteria have evolved discrete genetic systems that subvert this host-derived antimicrobial response. Systems for Cu and Zn homeostasis are well characterized, including the transcriptional control of sensing and responding to metal stress. Here we have discovered novel features of metal response sytems in *Streptococcus* that have major implications for pathogenesis and virulence. We show that *Streptococcus* resists Zn intoxication by utilizing a *bona fide* Cu regulator, CopY, to maintain cellular metal homeostasis, which enables the bacteria to survive stressful conditions. We identify new genes in *Streptococcus* that confer resistance to zinc intoxication, including several that have not previously been linked to metal ion homeostasis in any bacterium. The identification of cross-system metal management and new resistance mechanisms enhances our understanding of metal ion homeostasis in bacteria and its effect on pathogenesis.

## Introduction

In prokaryotic and eukaryotic cells, copper (Cu) and zinc (Zn) are important cofactors for metalloenzymes [1, 2]. When present in excess, however, Cu and Zn can cause cellular toxicity and, for example, can exert antimicrobial effects in subcellular areas within infected phagocytic cells [3, 4]. The double-edged sword of supporting cell physiology versus toxicity of Cu and Zn offers potential antimicrobial benefit for the control of bacterial pathogens and is of interest in studies of host-pathogen interactions [5–7]. On the one hand, Cu intoxication in bacteria can reflect enzyme inactivation, deregulation of metabolism, and/or redox stress, such as higher potential to generate reactive oxygen species [8]. Zn intoxication can reflect an ablation of uptake of essential manganese (Mn) [9], which can compromise the bacterial cell response to oxidative stress [10]; Zn can also disrupt central carbon metabolism [11]. Phagocytes such as macrophages and neutrophils can mobilise intracellular pools of Cu and Zn to pro-actively expose internalized bacteria to metal conditions that are antimicrobial [5, 12, 13]. In some pathogenic bacteria, this can be counteracted by activation of metal efflux mechanisms to thwart metal intoxication [14].

In bacteria, adaptation to metal excess and limitation is complex, but several defined systems are based on efflux proteins including P-type ATPases, which confer resistance to metal stress in different pathogens [1, 3]. In streptococci, discrete genetic systems for cellular management of Cu and Zn homeostasis act via the regulation of metal import and export machinery [9, 15, 16]; a system for Cu efflux utilizes the canonical *cop* operon, encompassing *copA* that encodes a ATPase efflux pump that extrudes cellular Cu ions, alongside a Cu-specific transcriptional regulator *copY*, that represses the operon [17, 18]. A system for Zn efflux uses a Zn-specific transcriptional response regulator, *sczA*, to control a Zn efflux transporter, encoded by *czcD* [15, 19]. These two systems of *copA-copY* and *czcD*-*sczA* for Cu and Zn export, respectively, have recently been characterized in group B *Streptococcus* (GBS), which responds to excess Cu and Zn by de- repressing *copA* via CopY to drive Cu export from the cell [20], and by activating *czcD* via SczA to regulate intracellular Zn levels [21], respectively. Molecular cross-talk between microbial Cu and Zn management systems can be proposed by several observations reported in prior studies of different pathogens. Zn was shown to bind CopY in *Enterococcus* [22], and was linked with a disruption in cellular Cu content in *Acinetobacter* [14]. In *Streptococcus pneumoniae*, Zn inhibits the expression of *copY*, which implies that Zn may act as a non-cognate co-represser of *copY* [17]. In *Pseudomonas stutzeri*, overlapping regulation of genes that mediate Cu and Zn resistance has recently been reported [34], suggesting that a core set of bacterial genes respond to Cu and Zn, and are co-regulated. A Cu-responsive regulatory system for Cu uptake in *Candida* was recently shown to encompass an Iron (Fe)-starvation stress response [23]. Collectively, these observations highlight the complexity of bacterial adaptation to metal excess and limitation, and also point to potential cross-talk mechanisms between metal management systems that might be used to support cellular homeostasis and effect bacterial fitness in distinct environments. However, a cross-talk mechanism of Zn-mediated signaling effects through *copY* as a means of bacterial adaptation to metal stress has not been defined.

We examined Zn management in GBS, as an opportunistic bacterial pathogen with defined metal detoxification systems in *copA-copY* and *czcD*-*sczA*, to determine whether CopY functions as a cross-system regulator of the bacteria’s response to Zn stress. We investigated the transcriptional links between the systems for Cu and Zn detoxification on a global scale, identified the complete genome of GBS that contributes to Zn resistance and examined the effects on bacterial virulence.

## Results

### Cross-over control of multiple metal efflux pathways by CopY

We recently defined the role of *copY* in regulating responses of GBS to Cu stress via control of *copA* [20], and *sczA* in regulating Zn stress via *czcD* [21]. Here, to examine cross-control of metal efflux pathways by non-cognate regulators, we compared the growth and metal stress resistance phenotypes of GBS mutants deficient in *copY* or *sczA* using defined *in vitro* conditions of either Cu stress (for *sczA*^-^) or Zn (for *copY*^-^) stress. This cross-comparative approach revealed unexpected cross-system regulatory effects of CopY towards the resistance of GBS to Zn intoxication. GBS was rendered severely susceptible to Zn intoxication as a result of *copY* mutation, according to growth analysis in a chemically-defined minimal medium (CDM) (Fig 1). *copY* also contributed to GBS resistance to Zn stress in nutritionally-rich growth conditions (THB medium) but was not significantly effected for growth in the absence of Zn stress (Supplementary Fig S1A and B). Together, these findings establish that *copY* exerts a key regulatory effect on the ability of GBS to respond to Zn intoxication.

**Figure 1.**
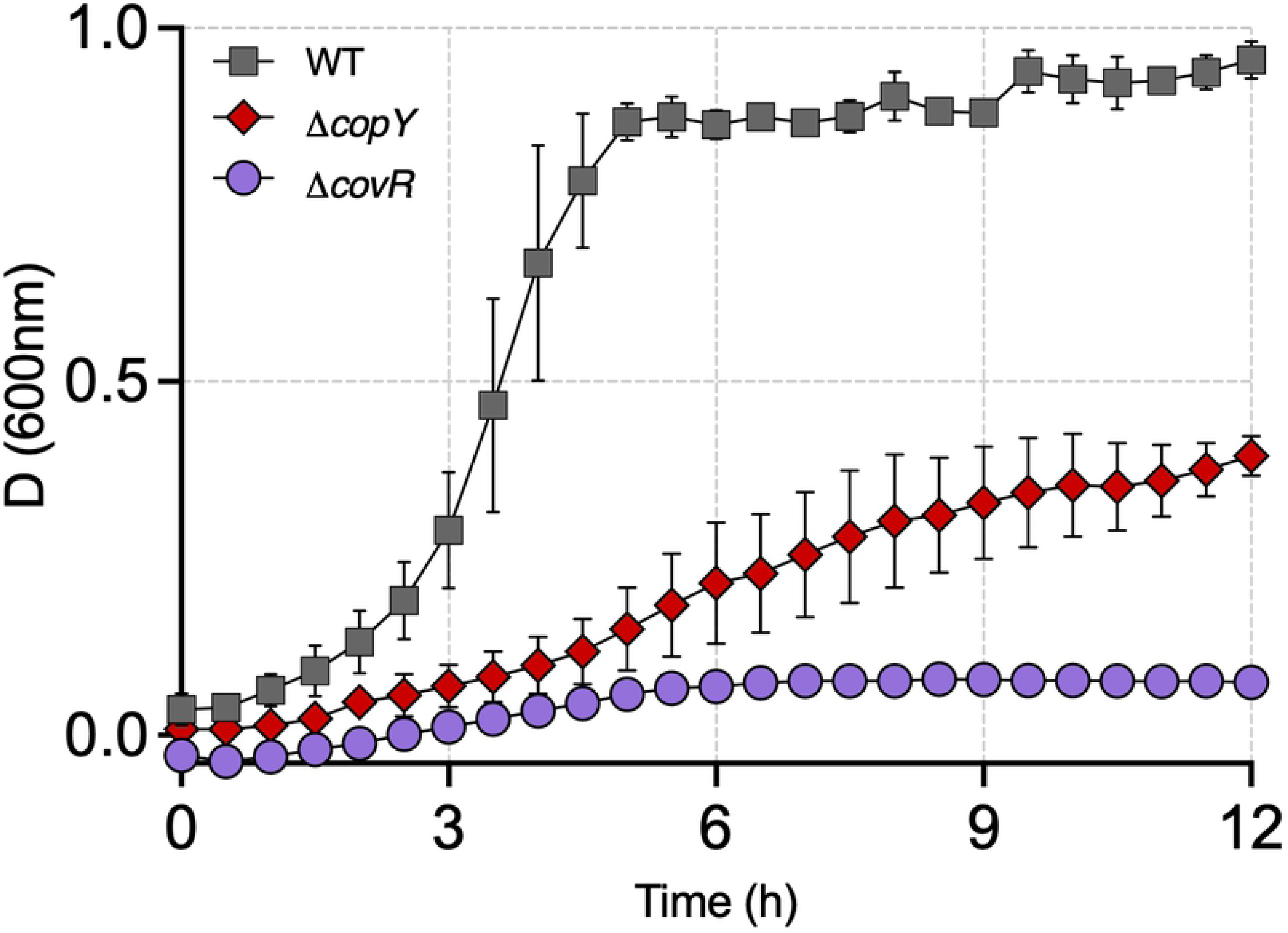
Growth analysis of GBS in rich and limiting medium and subjected to Zn intoxication. WT, *copY^-^* and *covR^-^* mutants were grown in nutrient-rich THB (A), THB medium supplemented with 1.5 mM Zn (B), nutrient-limited CDM (C) or CDM supplemented with 0.1 mM Zn (D) to examine responses to Zn stress. Bars show mean ± S.E.M (*n*=3 biological repeats).

Several pathogenic *Streptococcus* spp. respond to environmental stress cues, including excess metal ions (e.g. Mg^2+^) centrally via global response regulators, including *covRS* [24–26]. In GBS, a few genes that contribute to Zn homeostasis have also been linked to *covR*-regulation, with *adcR- adcCB* for Zn import in GBS strain 2603V/R, and *czcD* for Zn efflux in both 2603V/r and NEM316 strains [27, 28]. We therefore explored the cross-system effect of Zn and Cu stress at the point of *covR* by assessing the growth of *covR^-^* GBS in conditions of Zn or Cu stress. This showed a major contribution of *covR* in supporting GBS resistance to Zn intoxication (Fig 1) reflected in heightened susceptibility of the *covR* mutant to Zn stress. The mutant was also rendered more susceptible to Cu intoxication but to a lesser extent (Fig S1B and C). Thus, *covR* supports control of metal resistance in GBS in a manner that parallels the dual-metal resistance function of CopY towards Zn and Cu.

### CopY manages multiple intracellular metal pools during Zn stress

We used ICP-OES to measure the total intracellular content of metals in WT and *copY*^-^ GBS that were exposed to Zn stress, which showed that an absence of *copY* led to mis-management of the intracellular pools of multiple metals, including Zn, Cu, Mn, Fe and Mg (Fig 2). Comparing to WT in identical conditions, *copY*^-^ GBS had elevated levels of Zn, Cu, Mn, Fe and Mg within its cells. This broad level of mis-management of intracellular metal homeostasis in *copY*^-^ GBS is specific to Zn stress compared to Cu stress [20]. Thus, *copY*-mediated control of metal homeostasis in GBS extends beyond that of Cu, and enables the bacteria to manage the intracellular pools of multiple metals. Mutation in *covR* also effected the intracellular pool of some metals, including an elevation of Mn, and reduction in Fe levels compared to WT (Supplementary Fig S2). Thus, CopY broadly manages the intracellular pools of multiple metals in GBS exposed to non-cognate metal stress.

**Figure 2.**
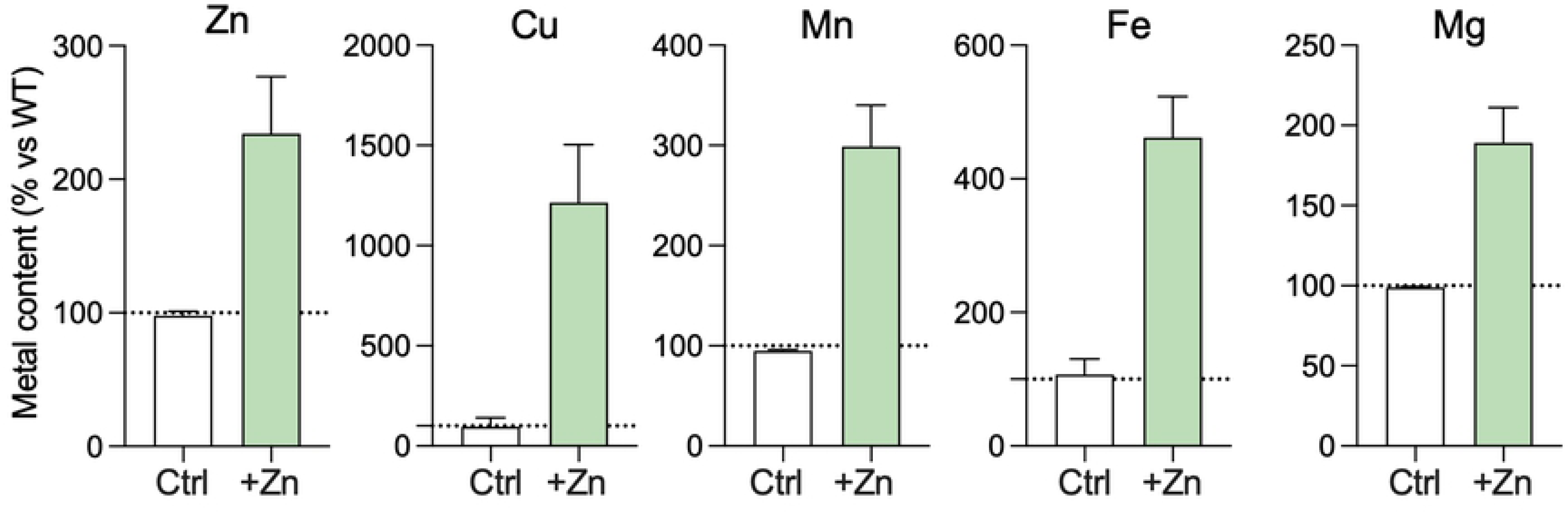
Intracellular accumulation of Zn, Cu, Mn, Fe or Mg in *copY*^-^ GBS with or without Zn stress. Cells of *copY*^-^ GBS were exposed to Zn (0.25 mM; green bars) and cellular metal content was compared to unexposed controls (THB only; white bars) using Inductively coupled plasma optical emission spectrometry (ICP-OES). Metal content (μg.g^-1^ dry weight biomass) normalised to WT cells from the same condition is shown as a percentage. Bars show mean ± S.E.M (*n*=3 biological repeats). Dotted line at 100% represents values equivalent to WT GBS.

### Transcriptional basis of CopY cross-system control of Zn homeostasis

To discern the effect of *copY* mutation on bacterial transcriptional responses to Zn stress, we analyzed genes that contribute to Zn resistance in GBS, namely *czcD* and *sczA*, using qPCR to measure gene expression in *copY*^-^ and WT GBS. Additionally, we analyzed Cu-responsive genes (*copA* and *copY*) in Zn stress, Cu stress and non-exposed controls. Unexpectedly, we detected a significant cross-system effect whereby *copY* was essential for the induction of *sczA* by Zn (Fig 3A). Interestingly, we also found that *copY* was controlled, in part, by *covR* in the response to Cu stress, but not Zn stess (Fig 3B); *i.e.,* Cu stress increased the quantity of *copY* mRNA in *covR^-^* GBS (versus WT in Cu stress). Similarly, activation of *sczA* in response to Zn stress did not occur in the *covR^-^* strain (Fig 3A). Notably, modulation of i) *sczA* by CopY (or CovR) did not effect *czcD* expression (Fig 3C), nor did modulation of *copY* by CovR affect *copA* expression (Fig 3D). This suggests that the sensitivity of *copY*^-^ GBS to Zn stress is not be explained by differences in Zn- efflux (*czcD* expression). Finally, *covR* contributes regulatory input as an auxiliary controller, to govern the expression of the regulatory genes that control Cu and Zn efflux as a master regulator.

**Figure 3.**
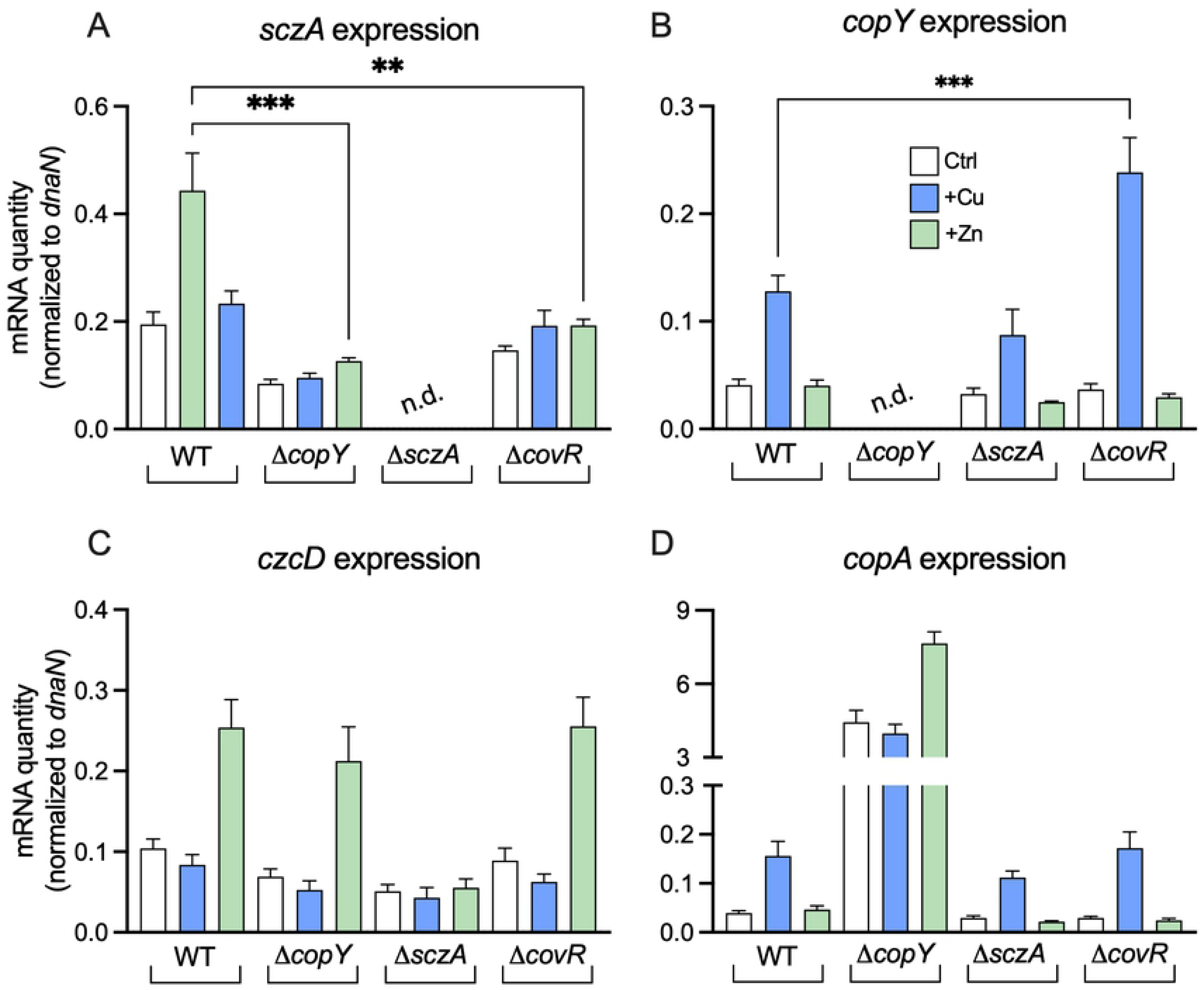
Cross-regulation of Zn and Cu stress responses by *copY* and *covR*. Transcripts of *sczA* (A), *copY* (B), *czcD* (C) and *copA* (D) were quantified by qRTPCR from cultures of WT, *copY^-^*, *covR^-^ and sczA^-^* mutants supplemented with Zn (0.25 mM) or Cu (0.5 mM) and compared to unexposed (THB only) controls (*n*=4). Absolute transcript amounts were normalized using housekeeping *dnaN* and generated from standard curves using GBS genomic DNA. Transcripts of *sczA* and *copY* were not detected (n.d.) in their respective mutant strains. Quantities were compared using ordinary one-way ANOVA and Holm-Sidak multiple comparisons. ** P < 0.01, ***P < 0.001.

### The copY-driven transcriptome of GBS exposed to Zn stress

We previously used RNA-seq to elucidate the transcriptome of GBS exposed to Zn stress, which identified >400 differentially expressed genes [21]. Here, we dissected the role of *copY* in the cell response to Zn stress using RNA-seq to compare *copY*^-^ GBS to WT exposed to Zn stress. The strains were grown in THB or THB supplemented with ± 0.25 mM Zn (not toxic for either strain) to facilitate a cross-strain comparison independent of any bias from growth-phase. In comparing transcriptomes of *copY*^-^ GBS to WT in the absence of Zn, in addition to massive de-repression of *copAZ* (∼200-fold up-regulated), we detected notable changes (±2-fold, P-adj < 0.05) that could partially explain the sensitivity of the *copY^-^* strain to Zn stress (Fig 4A). For example, we detected significant upregulation of *adcA* (2.3-fold) in *copY^-^* GBS, encoding a Zn import system in GBS [29], and a ∼4-fold reduction in expression of *arcABCD*, encoding an arginine deaminase system that confers a survival advantage under Zn stress [21]. Other dysregulated targets included genes for riboflavin synthesis (*ribDEAH* ∼10-fold down, see below) and transport (*ribU* ∼2-fold down), and a surface associated virulence factor (*fbsB* 2.5-fold up) (Dataset S1). In total, we identified 79 targets that were significantly altered in expression as a result of the loss of *copY* in GBS (see Dataset S1). Of note was a three-gene locus encoding *cyrR* (here, termed *<u>c</u>op<u>Y</u>*- *r*esponsive *<u>R</u>*egulator), *hly3* and *updK*, which was severely down-regulated (9 to 20-fold; (Fig 4A, Dataset S1); in *E. coli*, CyrR is part of the MerR superfamily that includes Zn- and Cu-responsive regulators ZntR and CueR; *hly3* encodes a putative hemolysin III/membrane protein but has not been investigated in GBS.

**Figure 4.**
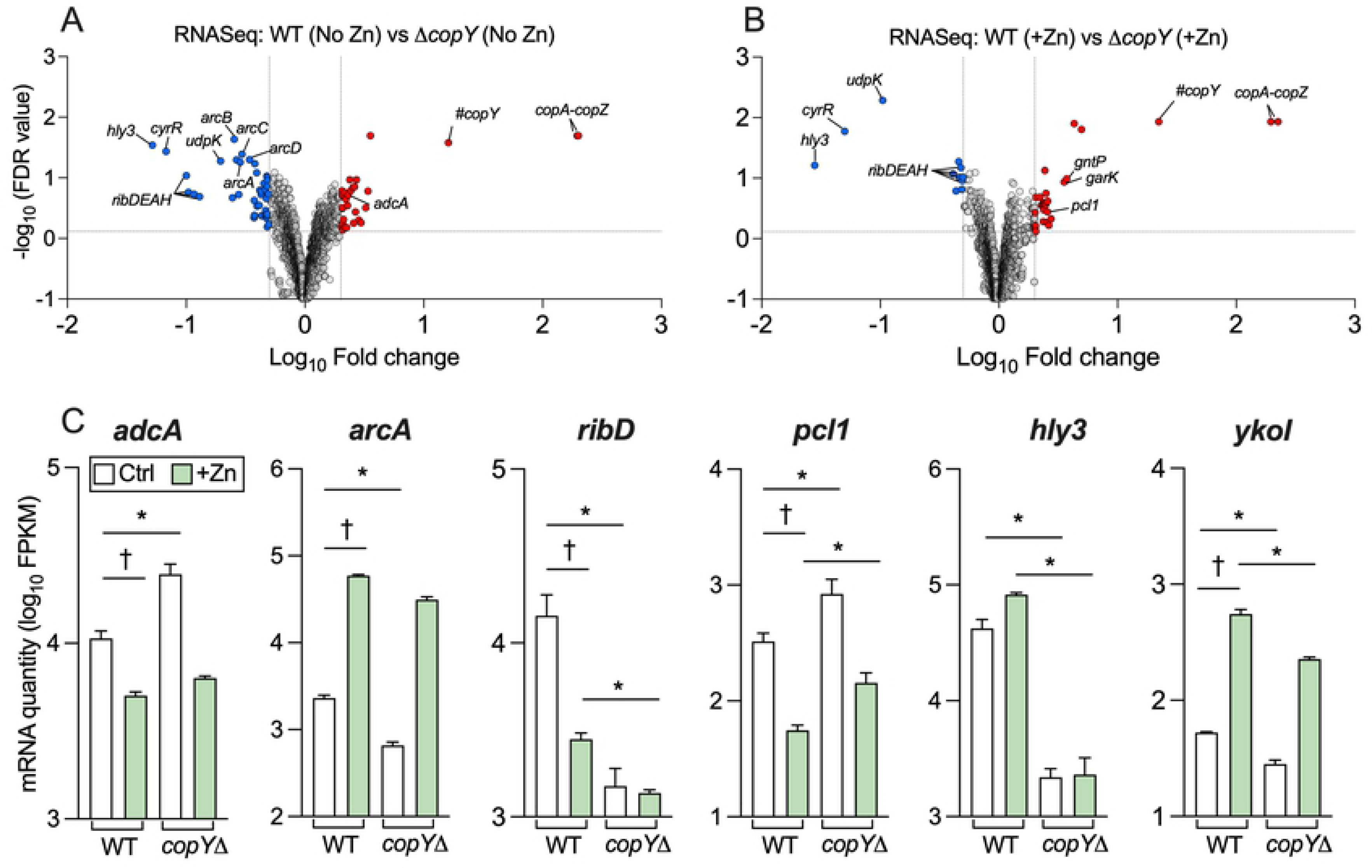
The CopY-responsive GBS transcriptome and the impact of Zn stress. Volcano plot showing data from RNASeq of WT GBS cultures compared to *copY-* GBS in THB (A), or THB supplemented with 0.25 mM Zn (B). Transcripts up- or down-regulated in response to Zn (n=4, >± 2-fold, FDR <0.05) are highlighted in red and blue, respectively. Dotted lines show False discovery rate (FDR; q-value) and fold change cut-offs. Grey points indicate genes that were unchanged. Selected genes are identified individually with black lines. Expression of individual genes from RNAseq analyses (C) showing mean Fragments Per Kilobase of transcript per Million mapped reads (FPKM) values for each condition in each strain. Data were compared with DESeq2 (* P-adj < 0.05 and ±2-fold; *n=*4). † indicates genes significantly altered in response to Zn stress in WT GBS as identified in previous work [21]. # reads mapped to truncated *copY* gene in Δ*copY* strain.

Next, we compared the transcriptomes of *copY*^-^ GBS and WT exposed to Zn stress, which revealed 78 transcripts that were significantly altered (±2-fold, P-adj < 0.05). Strikingly, although most transcript changes were shared between the strain comparisons independent of Zn stress (e.g. *cyrR-hly3*, *ribDEAH*, *pcl1*, *ykoI*), both *adcA* and *arcABCD* no longer responded significantly to Zn stress in the *copY^-^* strain (Fig 4B & C). Expression data for selected genes and conditions are shown in Fig 4C (complete set provided in Dataset S1).

Several genes identified by RNA-seq as being dysregulated were subsequently analyzed using qRTPCR to validate the responses of *copY^-^* GBS to Zn stress. We used WT GBS as a baseline to confirm selected Zn-dependent cross-system transcriptional effects that are controlled by CopY. This revealed five patterns of gene dysregulation, reflective of CopY cross-system effects, which are summarized in Fig 5; the patterns were Zn-induced genes that CopY (i) represses (*e.g. garK- gntP*) or (ii) activates (*e.g. ykoI*); (iii) Zn-repressed genes that CopY activates (*e.g. pcl1*), (iv) genes subject to Zn- and CopY-dependent de-repression (*e.g. ribDEAH*); and (v) genes activated by CopY irrespective of Zn (*e.g. cyrR-hly3-udpK*).

**Figure 5.**
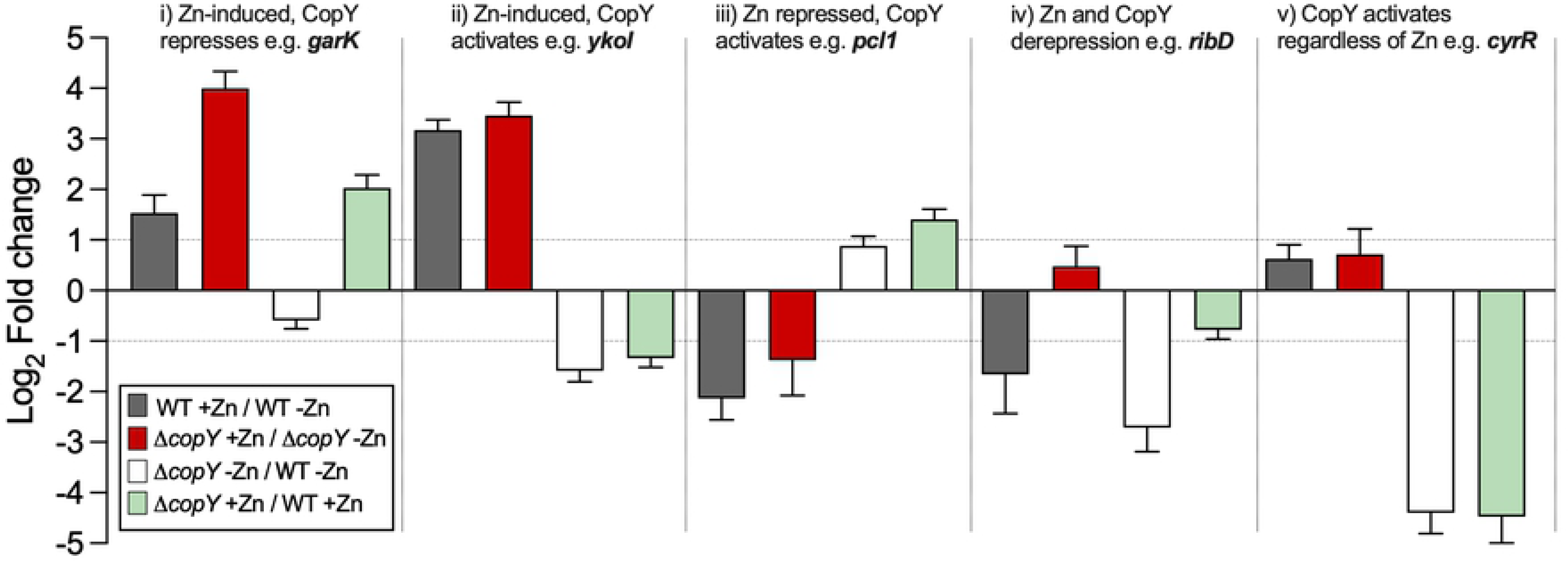
Cross system effects of CopY on Zn stress responses. Expression ratios (log_2_FC) of selected genes, identified by RNAseq as linked to CopY and Zn stress, were compared using qRTPCR as indicated, using RNA isolated from WT or *copY*^-^ GBS grown in THB or THB supplemented with 0.25 mM Zn (*n*=4). Five distinct activation states were apparent based on Zn and/or CopY dependency. Fold change values were calculated using *dnaN* as housekeeper and ΔΔ^CT^ values incorporated primer efficiency values as previously described [56].

The *ribDEAH* operon encodes a putative riboflavin synthesis pathway in GBS, and we found these genes were down-regulated in response to Zn stress in WT GBS in a prior study [21]. The down- regulation of these genes in response to *copY* mutation in this study hints at a connection between CopY and Zn stress that might effect bacterial metabolism. To functionally dissect the outcome of the transcriptional response of *ribDEAH* (considering its activation state was regulated by both Zn and CopY) we used an isogenic mutant in *ribD* and examined its phenotype in growth assays with Zn stress. This approach required the synthesis of a Modified Defined Medium (MDM; *Materials and Methods* and Supplementary Table S1) to examine growth in a defined medium deplete of riboflavin. In MDM lacking riboflavin, *ribD^-^* GBS did not grow and the *copY*^-^ strain grew poorly compared to WT (Fig S1A). In conditions of Zn stress (0.1mM), neither the *ribD^-^* nor the *copY*^-^ strain grew in MDM, however the growth of the WT was unaffected at this Zn concentration (Fig 6A). Supplementation of the media with 0.5 mg/L riboflavin and using the same Zn stress condition revealed that riboflavin restored growth to *ribD^-^* GBS, however *copY^-^* GBS exhibited a severe attenuated phenotype in this condition (Fig 6B). Growth of WT GBS was unaffected in the absence of riboflavin, consistent with a functional *ribDEAH* operon and confirming a role for these genes in *de novo* synthesis of this vitamin; all three strains grew in MDM supplemented with riboflavin (but without Zn; Fig S3). *ribD* and several other targets subject to CopY regulation (e.g. *pcl1*, *ykoI* and *hvgA*) were likely co-regulated by CovR because their transcription was altered comparing WT to *covR^-^* GBS (Fig S4). Thus, *copY* plays a central role the GBS Zn stress response by regulating gene targets at the transcriptional level; genes regulated by *copY* cued by Zn stress effect GBS growth capacity. Transcriptional co-regulation of Cu and Zn export responses by *covR* provides auxiliary control beyond *copY* to manage metal stress in GBS.

**Figure 6.**
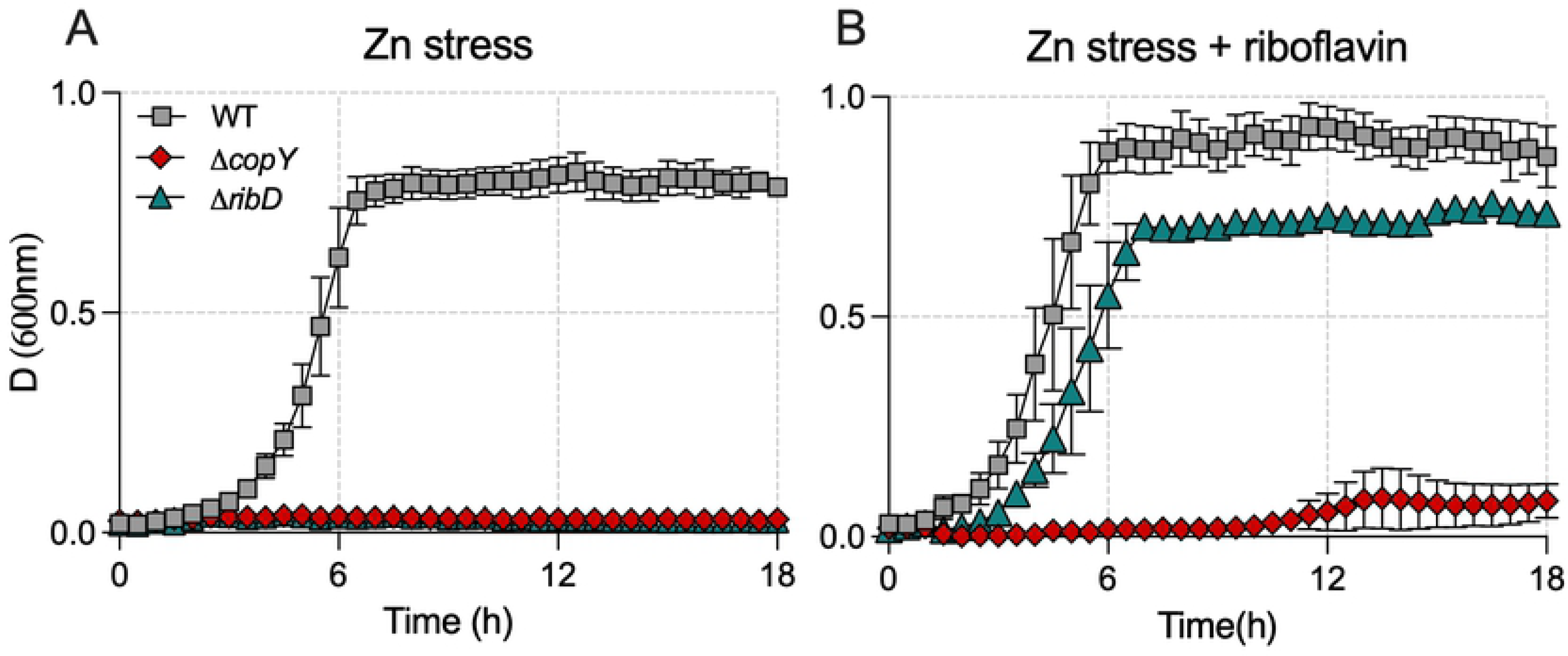
The effect of riboflavin on Zn stress resistance in GBS. WT, *copY^-^* and *ribD^-^* mutants were grown in MDM supplemented with 0.1 mM Zn (A) or MDM supplemented with 0.1 mM Zn and 0.5 mg/L riboflavin (B). Bars show mean ± S.E.M (*n*=3 biological repeats) measures of attenuance (*D* at 600_nm_).

To analyze an additional *copY*-regulated target cued by Zn stress in GBS and test the functional outcome of bacterial growth, we mutated *hly3* and analysed the growth of the *hly3^-^* strain with and without Zn stress in both nutritive and nutrient-limited medium. We found the isogenic *hly3^-^* strain was significantly impaired for growth in THB (Fig. S5A) but this attenuation was absent from comparisons of WT to *hly3^-^* mutant in Zn stress conditions in THB (Fig S5B) or in CDM (Fig S5C and D). These data suggest that *hly3* contributes to growth activities that occur in THB, but not CDM, that are disrupted during Zn stress. These findings establish that *hly3* is a target of *copY* in response to Zn stress, and it likely contributes to growth of GBS in certain conditions. Together, these findings demonstrate that GBS engages CopY in response to multiple metal stress cues to enable a coordinated gene expression response that extends beyond the *cop* operon to support bacterial survival during metal stress.

### CopY is conserved among streptococci and supports virulence

The function of *copY* that confers a transcriptional mechanism of cross-system control to respond to Zn stress in GBS prompted us to examine conservation among other pathogenic streptococci, and ascertain whether it contributes to pathogenesis. Sequence analysis and structural modelling revealed that *copY* is highly conserved among multiple GBS strains, as well as other pathogenic *Streptococcus* spp., and other pathogenic gram-positive cocci (Fig 7). This modelling enabled a schematic representation of the putative metal binding site at the C-terminus of the *S. agalactiae* CopY (Fig 7). In the absence of any ascribed role for *copY* in the pathogenicity of any bacterium, we tested whether *copY* contributes to GBS virulence using a model of systemic disseminated infection. Remarkably, *in vivo* infection assays in mice showed that *copY* was critical to GBS virulence; we observed that *copy^-^* GBS was severely attenuated in the blood, heart, lungs, spleen and kidneys of mice at 24h post-infection (Fig 8). Expectedly, *covR^-^* GBS was also attenuated in the blood, heart and lungs but was recovered in higher numbers from brain and liver compared to WT (Fig 8), pointing to tissue-specific effects. Thus, *copY* functions to support bacterial virulence.

**Figure 7.**
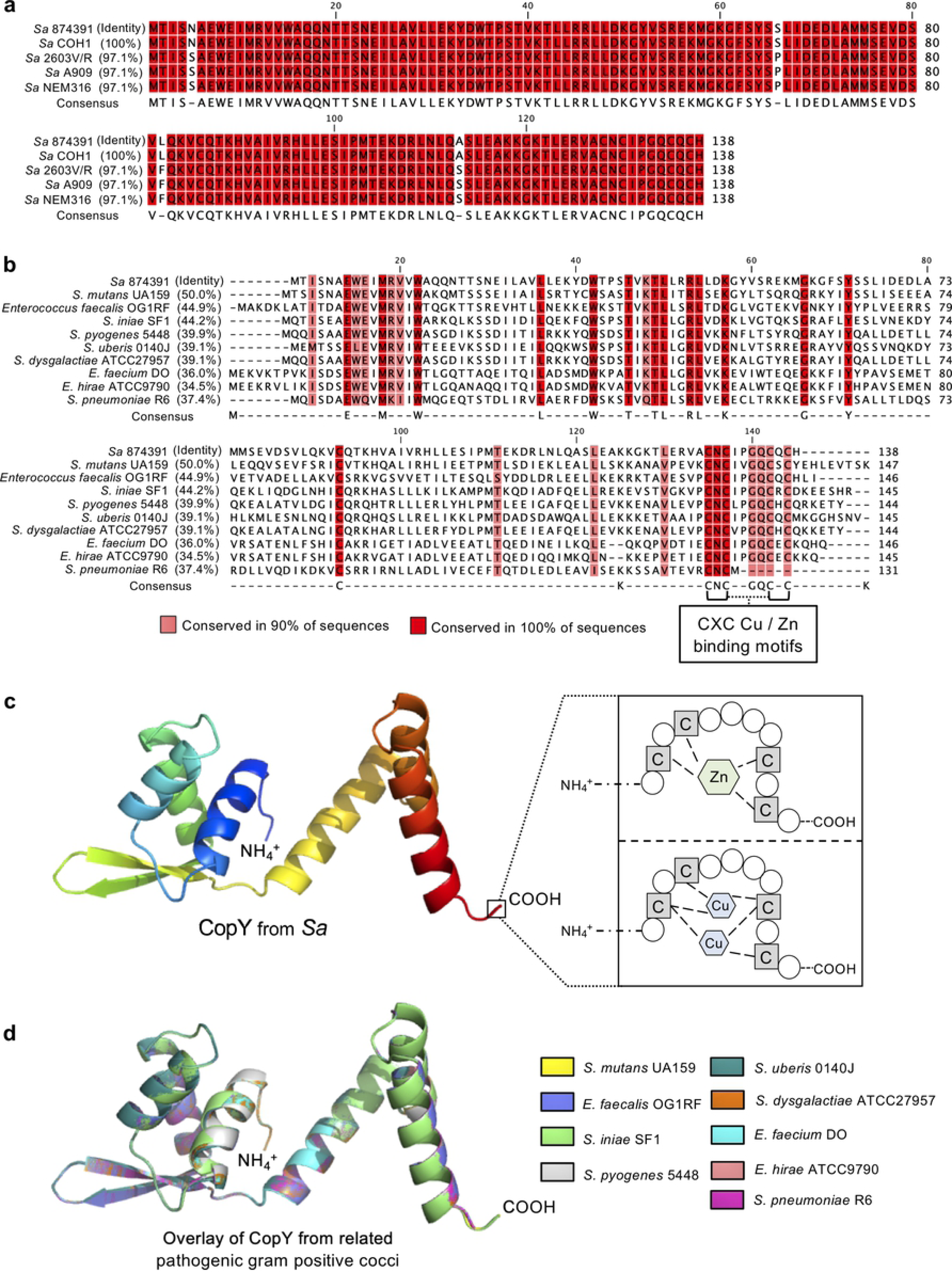
CopY likely operates as a cross-system regulator in numerous gram-positive bacteria. Alignment of CopY shows a high degree of conservation (>97% identity between reference *S. agalactiae* strains) (a). Alignment of *S. agalactiae* CopY with other *Streptococcus* and *Enterococcus* strains. Highlighted are two conserved putative CXC motifs that are predicted to bind Cu and/or Zn at the C-terminus; amino acids that are >90% conserved are shaded in red, as indicated (b). Predicted structural model of *S. agalactiae* CopY and schematic representation of the putative metal binding site at the C-terminus of the protein, adapted from (Cobine *et al*., 2002) (c). Structural alignments of predicted CopY proteins from *Streptococcus* and *Enterococcus* strains indicate overlapping protein conformation despite modest conservation of amino acid identity (d).

**Figure 8.**
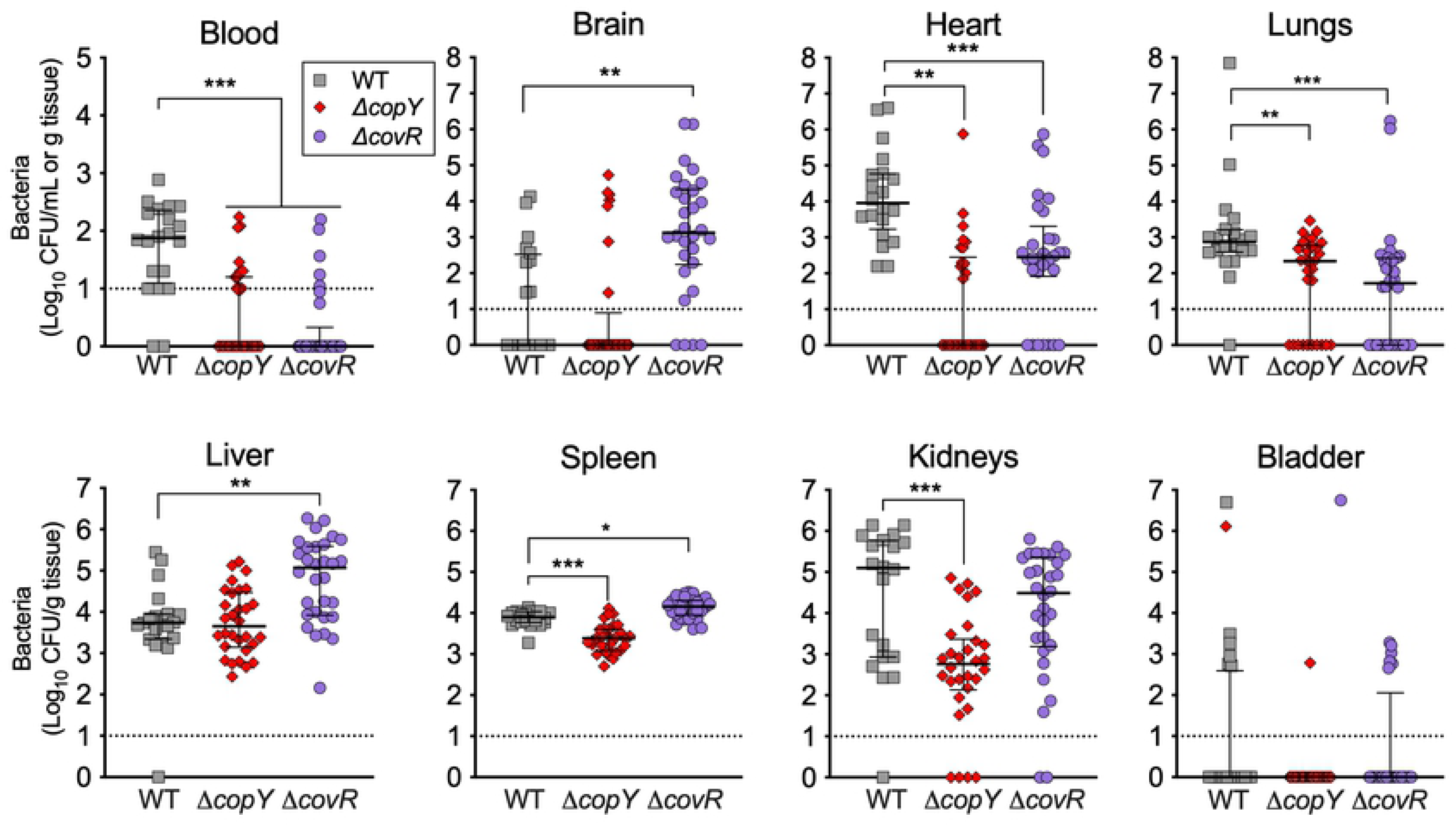
Mutation in *copY* and *covR* have major implications in colonization and disseminated spread of GBS bloodstream infection. Virulence of WT (grey squares), Δ*copY* (red diamonds) or Δ*covR* GBS (purple circles) in a mouse model of disseminated infection. C57BL/6 mice (6-8 weeks old) were intravenously injected with 10^7^ bacteria; bacteremia and disseminated spread of bacteria to brain, heart, lungs, liver, spleen, kidneys and bladder were monitored at 24h post infection. CFU were enumerated and counts were normalized using tissue mass in g. Viable Cell counts of 0 CFU/mL were assigned a value of 1 to enable visualisation on log_10_ y-axes. Lines and bars show median and interquartile ranges and data are pooled from 2-3 independent experiments each containing n=10 mice; groups infected with mutants were compared to WT group using Kruskal-Wallis ANOVA with Dunn’s corrections for multiple comparisons (*P < 0.05, **P < 0.01, *** P < 0.001).

### Forward genetic screen for mediators of GBS resistance to Zn stress

To examine the entire GBS genome for functionally related regions that contribute to resistance to Zn stress we used an open-ended approach based on a super-saturated ∼430,000-mutant library, generated using pGh9-IS*S1* [30]. We exposed the bacteria to Zn stress, comparing to non-exposed controls *en masse*. Stringent selection criteria (±4-fold, P-adj < 0.05) identified 12 genes that were essential for GBS to survive during Zn stress; insertional site mapping revealed the frequency of insertions was significantly under-represented in these 12 genes (Fig 9A and Dataset S2). Conversely, 26 genes for which the mapped insertions were over-represented were identified, suggesting that constrain GBS growth in Zn stress. Representative mapping is shown for selected genes in Fig 9B-D. To validate these hits, we generated targeted isogenic mutants of several candidate genes of the Zn stress resistome, including *stp1* and *stk1* (CHF17_00435 and CHF00436; serine/threonine phosphatase and kinase pair), *celB* (CHF17_01596; EIIC disaccharide transporter), *rfaB* (CHF17_00838; glycosyltransferase), and *yceG* (CHF17_01646; *mltG*-like endolytic transglycosylase). In comparing the growth of WT GBS to mutants in Zn, we detected attenuation in all mutants (for under-represented genes) (Fig 10A-B). Notably, some strains exhibited growth defects in the absence of Zn (e.g. Δ*stp1,* Δ*stk1* and Δ*plyB*). Mutation of *arcR* (over-represented) showed a hyper-resistance phenotype; the Δ*arcR* mutant grew in high Zn (CDM with 0.25 mM; Fig 9C), which approached inhibitory for the WT (Fig 10A). Together, these findings identify a suite of genes in the GBS genome that contribute to the bacteria’s ability to resist Zn intoxication, and which function either by supporting or constraining growth of GBS.

**Figure 9.**
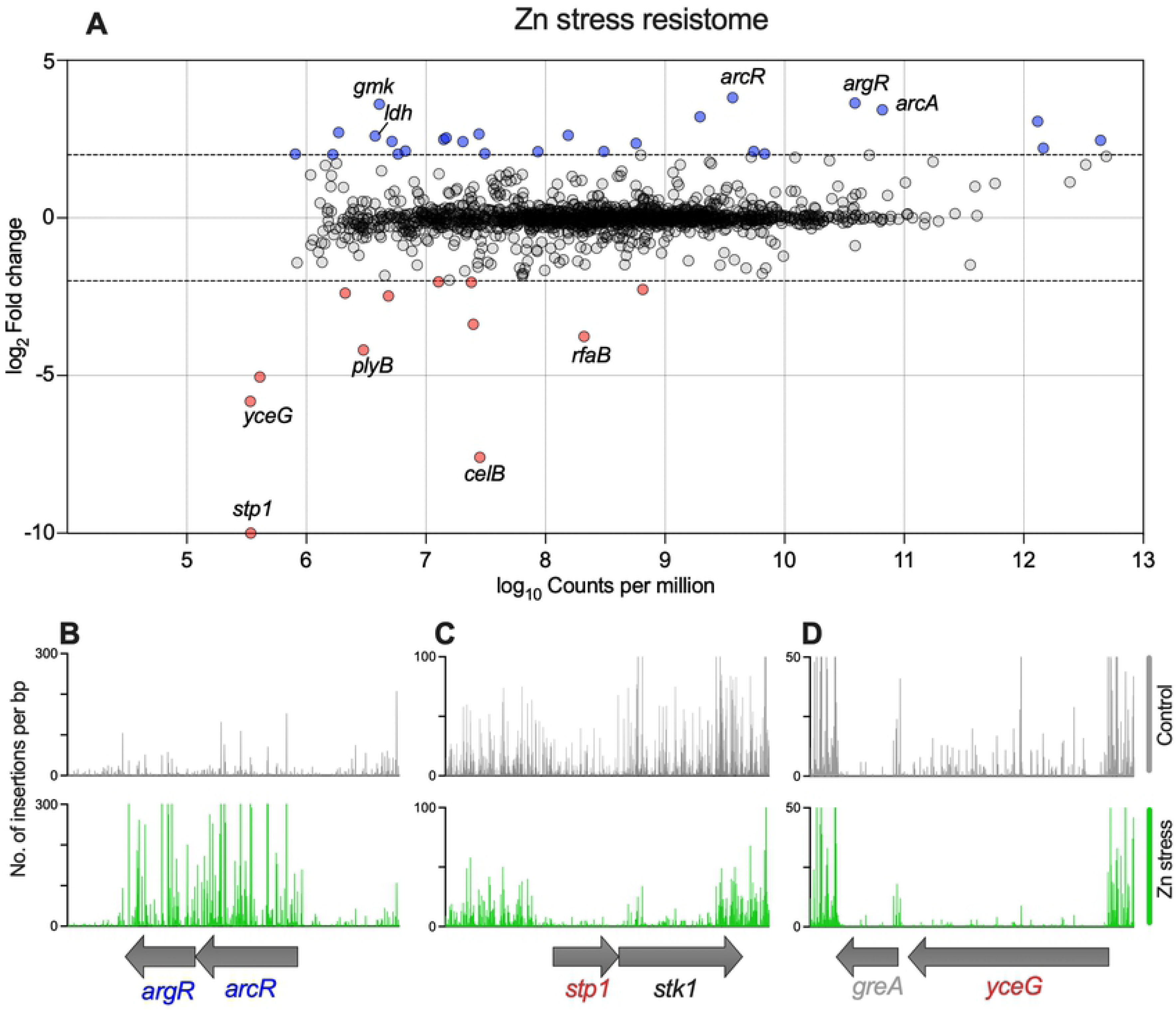
Defining the Zn stress resistome of *S. agalactiae* to identify novel factors in bacterial responses to Zn intoxication. A super-saturated IS*S1 S. agalactiae* insertion library was subjected to Zn stress and compared to control incubation without Zn to define the Zn resistome (A). Transposon-directed insertion sequencing (TraDIS) identified 26 genes over- represented (blue) and 12 under-represented (red) during Zn stress (cutoffs: FDR < 0.05, fold- change ±4). Illustrative read-mapping of IS*S1* insertion sites (B-D) displaying differences between non-exposed control (grey) or Zn stress conditions (green) for selected genes; over-represented *argR/arcR* (B) and under-represented *stp1/stk1* (C) or *yceG* (D). Vertical lines in B-D represent pooled read counts at each base within locus, with coding sequences of genes represented by grey arrows beneath. Data are compiled from 3 independent experiments.

**Figure 10.**
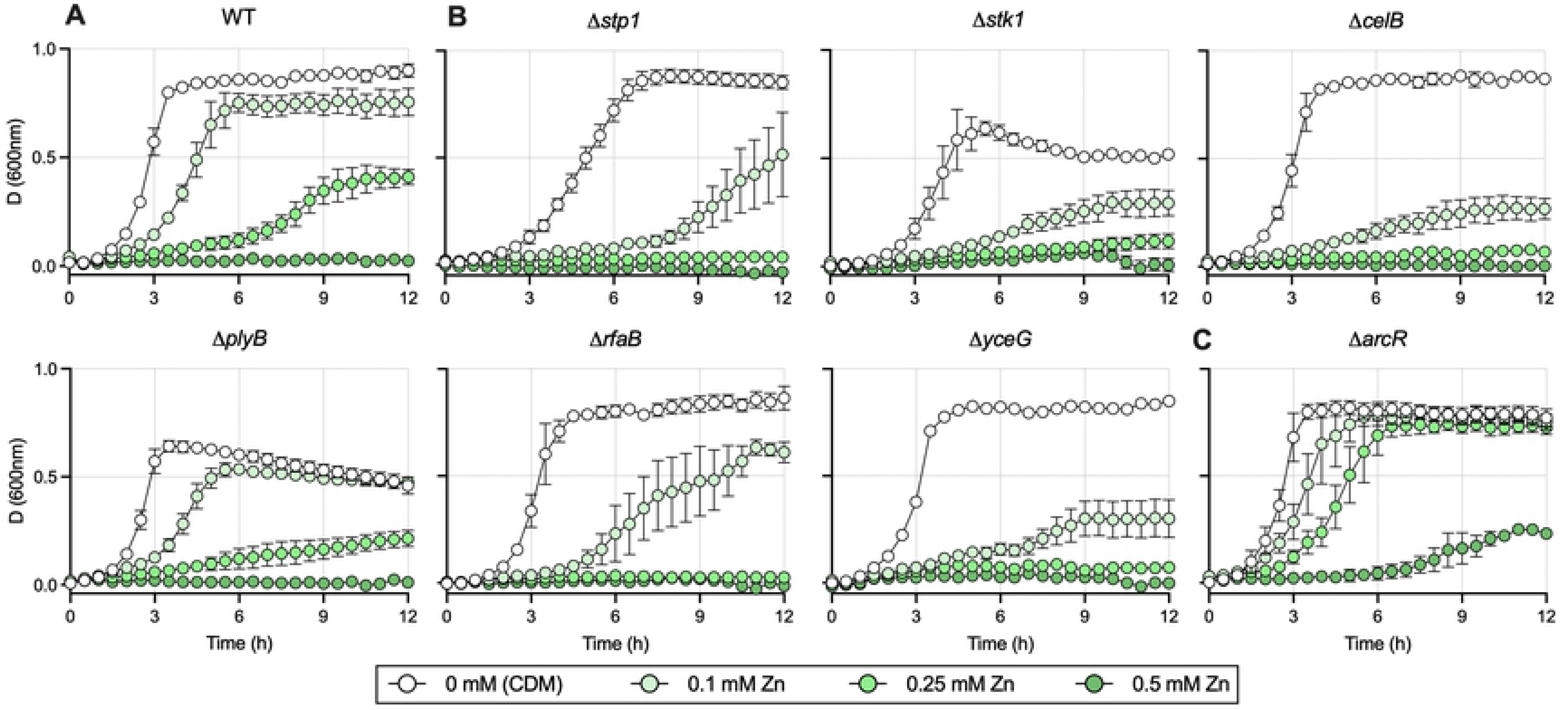
Validation of TraDIS hits by isogenic mutation and phenotypic comparisons for selected genes of the Zn stress resistomes. WT *S. agalactiae* (A) and mutants with deletions in genes identified as important during in Zn intoxication by TraDIS (B-C) were examined for growth phenotypes in CDM (a nutrient-limited medium) or CDM supplemented with 0.1, 0.25 and 0.5mM Zn as indicated. Points show means of attenuance (600nm) and bars show s.e.m. (*n*≥3).

## Discussion

One important function of metalloregulatory proteins is to bind and respond to cognate effectors, while ignoring non-cognate (competing) metals. In bacteria, this functional feature facilitates co- ordinated expression of metal acquisition systems in conditions of metal limitation, whereas, during metal excess, it enables bacteria to drive efflux systems to underpin divergent, contrary responses and resist metal intoxication [3]. The dogma of the function of the CopY transcriptional regulator in bacteria has until now centred on the management of Cu homeostasis via direct effects on the *cop* operon, in response to cues from its cognate effector, Cu. Despite the essential role of CopY in Cu homeostasis in bacteria, a hypothesis that CopY is nonresponsive to competing metals has not directly been addressed until now. This study establishes a new, biologically consequential function of CopY is the mediation of cellular responses to Zn stress in bacteria, which for GBS entails (i) a fundamental role of *copY* in conferring bacterial resistance to Zn intoxication, (ii) robust regulatory inputs from *copY* in response to Zn stress cues, which drive cross-system effects to support bacterial Zn homeostasis, (iii) a virulence function of *copY* that promotes GBS survival in acute disseminated infection, and (v) a defined 46-member family of targets in GBS that comprise the Zn resistome. Overall, this study shows that *copY* controls two discrete systems for Cu and Zn homeostasis in *Streptococcus*, and establishes a collection of genomic elements that enable the bacteria to survive Zn intoxication.

The transcriptional landscape of GBS in response to Zn defined here elucidates a CopY-regulon of Zn-responsive targets, which represents the first described in a bacterial pathogen. In defining the cross-system effects of CopY that stem from exposure to Zn stress, this study reveals that *copY* induces robust expression of all the genes that make up the *cop* operon. *copY* regulates a small but distinctive group of additional targets, including multiple genes that have no known links with metal stress responses in bacteria nor virulence (e.g., *hly3, cyrR, ribD, ykoI, garK*). One of these, CyrR, is of the MerR superfamily that includes Zn- and Cu-responsive regulators ZntR and CueR of *E. coli* [1, 31]. Identification of a three-gene locus of *cyrR*, *hly3* and *updK*, which GBS down-regulates in response to Zn stress supports the hypothesis that this locus responds to various regulatory inputs, as reported previously [32, 33]. Intruigingly, in our study, this response required intact *copY*, revealing a novel mechanism of transcriptional control of the *cyrR*-*hly3*- *updK* locus. Another locus of the CopY-regulon of Zn-responsive targets, *ribDEAH*, supports riboflavin biosynthesis; *ribD* has no prior known links with Zn or Cu resistance in bacteria. Testing of a targeted mutant for *ribD* revealed a contribution of riboflavin synthesis to resisting Zn stress. Riboflavin supports an array of metabolic processes because it’s downstream products are flavin coenzymes, flavin mononucleotide (FMN) and flavin adenine dinucleotide (FAD), as required for oxidative metabolism and other processes; in addition, FMN can act as a precursor to cobalamin synthesis [34]. GBS contains a putative homologue of a riboflavin import protein RibU (ASZ01710.1) which is down-regulated ∼2-fold in the *copY*^-^ background (Dataset S1). In *S. pyogenes*, well studied for Zn intoxication resistance [11, 15, 35], the *ribDEAH* genes are absent, such that the organism relies solely on riboflavin import [36]. In *S. pneumoniae*, differential expression of *ribDEAH* leads to differences in host responses to different clinical isolates [37]. Precisely how riboflavin synthesis, versus uptake, (in bacteria such as GBS that can do both) contributes to attenuation of growth during Zn stress will need to be examined in future work.

In analyzing the function of *copY* compared to the Zn responsive regulator *sczA*, this study reveals the opposing nature of these two metalloregulatory proteins. The former responds to non-cognate Zn cues, but the latter is essentially non-responsive to Cu. We demonstrate that GBS in Zn stress utilizes CopY to regulate the intracellular pools of multiple metals in addition to Cu. This supports a model in which the ability of CopY to mediate cross-system effects is specific and functionally distinct from another key metalloregulatory protein. Structural modelling shows a high degree of conservation of CopY among pathogens closely related to GBS, which implies that cross-system effects for management of responses to metal stress at the point of *copY* may operate in other bacteria. These conserved aspects of CopY among related bacteria suggest that there will be utility in testing the function of *copY* in response to non-cognate metal stress in other bacteria.

The TraDIS in this study provides a comprehensive analysis of the Zn stress resistome of GBS, and identifies multiple targets new to the bacterial metal detoxification field [38]. Several identified as being most strongly associated with GBS survival in Zn stress (e.g. *arcR, rfaB, plyB, yceG, celB*, *stp1*) have not previously been linked to metal stress responses in any bacteria. TraDIS was used recently to study GBS survival in blood revealed effects of calprotectin [39–41]. The suite of genes encoding regulators and putative effectors that confer GBS resistance to Zn stress identified in this study dramatically expands our understanding of metal management in bacteria by offering new insight into the diversity of genes that mediate resistance to Zn intoxication, such as those encoding enzymes for metabolism and cell wall synthesis, transporters, and global transcriptional regulators. Our approach identified *arcR* and *argR,* two adjacent regulators that likely co-ordinate arginine deaminase (encoded by *arcABC*) expression, as constraining Zn resistance in GBS, since IS*S1* insertions were significantly enriched in *arcR* and *argR*. Isogenic mutation in *arcR* enhanced resistance to Zn, consistent with the TraDIS finding. A previous study identified a role for *arcA* in conferring resistance to Zn stress, since an isogenic *arcA* mutation attenuated growth under Zn intoxication conditions [21]. Interestingly, in contrast to this observation, we detected enrichment, rather than reduction, in IS*S1* insertions in *arcA*. This could be explained by a potential for polar effects of IS*S1* insertion on the *arcABDC* locus. It would be of interest to examine the contribution of *arcBC* and *arcD* to Zn resistance, since these encode proteins that produce or import ornithine, which was recently shown to rescue Zn sensitivity in GBS [21].

Analysis of CovR/CovS in the GBS response to Zn stress revealed a role for the Stk1/Stp1-CovR regulation axis in mediating Zn resistance. Stp1/Stk1 phosphorylate CovR to drive its effects [42, 43] and we found *stp1/stk1* were essential for Zn resistance. Together, these findings show that Stk1/Stp1-CovR regulatory activity helps to support Zn resistance in GBS. The CovR/CovS two-component system has been linked to streptococcal virulence but has not previously been linked with a response to Zn stress. That *covR* promotes resistance to Zn intoxication in GBS can be used to suggest a parallel between the *covRS* system and the dual-metal resistance regulatory function of *copY*, whereby the regulator governs resistance of the bacteria to multiple metals. A two-component system in *Caulobacter crescentus*, UzcRS, is highly responsive to both Zn and Cu (and uranium) to couple a response regulator to different extracytoplasmic metal stress responses [44]. In *Pseudomonas stutzeri*, overlapping regulation for Cu and Zn resistance genes was recently reported [45], with cross-regulation achieved by a core set of *P. stutzeri* Cu and Zn-responsive genes. In *Mycobacterium tuberculosis*, two paralogous ATPases, CtpD and CtpJ that are activated by Co(2+) and Ni(2+) appear to mediate metal efflux, but play non-redundant roles in virulence and metal efflux [46]. Further elucidation of cross-talk mediated by CopY, and CovR as a regulator governing resistance of GBS to multiple metals will be help to more clearly define their functions in comparison to systems in other bacterial pathogens.

Bacterial resistance to metal stress is used by some pathogens to evade host defences [12, 47]. Cu management contributes to virulence in some infections, but a role for CopY in virulence has not been reported. For example, *S. pneumoniae* regulates central metabolism in response to metal stress to support bacterial survival [41], and uses CopA to drive virulence during host infection [17]. In *E. coli*, Cu-transporting ATPases, including CopA are required for bacteria survival in an *in vitro* host-pathogen interface in macrophages [48]. We found that *copY* contributes to the virulence of GBS because *copY^-^* GBS was attenuated in multiple organs, including the blood, heart, lungs, and kidneys of mice following systemic infection. These findings are consistent with prior observations that have alluded to a role of CopY in supporting bacterial virulence. For example, increased expression of *copY* in *S. pneumoniae* in the lungs of mice was reported [17], and higher Cu levels along with co-incidental up-regulation of *copYAZ* in the blood of mice infected with *S. pyogenes* was reported [16]. Our findings for the *covR* mutant show that this global virulence regulator supports *S. agalactiae* survival but this depends on tissue context; the role of CovR in brain infection is consistent with a prior report [43].

The finding of massive upregulation (200-fold) of *copA-copZ* in *copY^-^* GBS is notable because this is based on a non-polar, unmarked deletion that we generated in this study. It would be of interest to determine if such transcription is translated into an enhancement of CopA-CopZ proteins of this magnitude. If so, this would place a significant metabolic burden on RNA and protein synthesis machinery and presumably might attenuate GBS growth. Differential insertion of IS*S1* was not detected in *copY* under Zn stress in the TraDIS analysis. This could be accounted for by the potential for polar effects of IS*S1* insertion in *cis,* thus abolishing, rather than up- regulating, *copA-copZ* transcription. Pointedly though, although growth rate of the *copY^-^* strain was reduced in THB medium conditioned for Zn stress (Fig 1B), this strain was able to achieve significant culture densities after 12h of growth. In support of this, several of the novel Zn resistome targets (e.g. *rfaB*, *stp1*, *celB*) exhibited a more dramatic attenuation phenotype compared to the *copY*^-^ strain (Fig S6). Future studies might utilise a nutrient-limited medium such as CDM or MDM with TraDIS and Zn stress to yield new factors that support GBS Zn resistance. In summary this study identifies a new role of *copY* in responding to Zn stress in GBS, revealing novel regulatory cross-talk between this Cu-sensing repressor that results in modulation of Zn homeostasis. The Zn sensitivity phenotype of *copY*-deficient GBS is not attributed to a single Zn-resistance effector (such as the CzcD efflux system) but arises from pleiotropic effects that encompass multiple factors that underpin GBS survival during Zn stress. These include arginine deaminase expression (via *arcABDC*), Zn import (via *adcA*) riboflavin synthesis (via *ribDEAH*) and an as-yet undefined role for the *cyrR-hly3* locus.

## Materials and Methods

### Bacterial strains, plasmids and growth conditions

GBS, *E. coli* and plasmids used are listed in Supplementary Table S2. GBS was routinely grown in Todd-Hewitt Broth (THB) or on TH agar (1.5% w/v). *E. coli* was grown in Lysogeny Broth (LB) or on LB agar. Routine retrospective colony counts were performed by plating dilutions of bacteria on tryptone soya agar containing 5% defibrinated horse blood (Thermo Fisher Scientific). Media were supplemented with antibiotics (spectinomycin (Sp) 100μg/mL; chloramphenicol (Cm) 10 μg/mL), as indicated. Growth assays used 200μL culture volumes in 96-well plates (Greiner) sealed using Breathe-Easy® membranes (Sigma-Aldrich) and measured attenuance (*D*, at 600nm) using a ClarioSTAR multimode plate reader (BMG Labtech) in Well Scan mode using a 3mm 5x5 scan matrix with 5 flashes per scan point and path length correction of 5.88mm, with agitation at 300rpm and recordings taken every 30min. Media for growth assays were THB and a modified Chemically-Defined Medium (CDM) [29] (with 1g/L glucose, 0.11g/L pyruvate and 50mg/L L-cysteine), or Modified Defined Medium (MDM; see Supplementary Table S1) supplemented with Cu or Zn (supplied as CuSO_4_ or ZnSO_4_) as indicated. For attenuance baseline correction, control wells without bacteria were included for Cu or Zn in media alone.

### DNA extraction and genetic modification of GBS

Plasmid DNA was isolated using miniprep kits (QIAGEN), with modifications for GBS as described elsewhere [49]. Mutant strains (Supplementary Table S2) were generated by isogenic gene-deletions, constructed by markerless allelic exchange using pHY304aad9 as described previously [21, 50]. Plasmids and primers are listed in Supplementary Table S2 and Supplementary Table S3, respectively. Mutants were validated by PCR using primers external to the mutation site and DNA sequencing.

### RNA extraction, qRTPCR

For Cu and Zn exposure experiments, 1mL of overnight THB cultures were back-diluted 1/100 in 100mL of THB (prewarmed at 37°C in 250mL Erlenmeyer flasks) supplemented with 0.25 mM Zn or 0.5 mM Cu. Cultures were grown shaking (200rpm) at 37°C; after exactly 2.5h, 10-50mL volumes containing approximately 500 million mid-log bacteria were harvested; RNA was preserved and isolated as described previously [51]. RNA quality was analysed by RNA LabChip using GX Touch (Perkin Elmer). RNA (1000ng) was reverse-transcribed using Superscript IV according to manufacturer’s instructions (Life Technologies) and cDNA was diluted 1:50 in water prior to qPCR. Primers (Supplementary Table S3) were designed using Primer3 Plus [52, 53] to quantify transcripts using Universal SYBR Green Supermix (Bio-Rad) using a Quantstudio 6 Flex (Applied Biosystems) system in accordance with MIQE guidelines [54]. Standard curves were generated using five-point serial dilutions of genomic DNA (5-fold) from WT GBS 874391 [55]. Expression ratios were calculated using C_T_ values and primer efficiencies as described elsewhere [56] using *dnaN*, encoding DNA polymerase III β-subunit as housekeeper.

### Whole bacterial cell metal content determination

Metal content in cells was determined as described [10]. Cultures were prepared essentially as described for *RNA extraction, qRTPCR* with the following modifications; THB medium was supplemented with 0.25 mM Zn or 0.5 mM Cu or not supplemented (Ctrl), and following exposure for 2.5h, bacteria were harvested by centrifugation at 4122 x g at 4°C. Cell pellets were washed 3 times in PBS + 5mM EDTA to remove extracellular metals, followed by 3 washes in PBS. Pelleted cells were dried overnight at 80°C and resuspended in 1mL of 32.5% nitric acid and incubated at 95°C for 1h. The metal ion containing supernatant was collected by centrifugation (14,000 x g, 30min) and diluted to a final concentration of 3.25% nitric acid for metal content determination using inductively coupled plasma optical emission spectroscopy (ICP-OES). ICP-OES was carried out on an Agilent 720 ICP-OES with axial torch, OneNeb concentric nebulizer and Agilent single pass glass cyclone spray chamber. The power was 1.4kW with 0.75L/min nebulizer gas, 15L/min plasma gas and 1.5L/min auxiliary gas flow. Cu was analysed at 324.75nm, Zn at 213.85nm, Fe at 259.94nm and Mn at 257.61nm with detection limits at <1.1ppm. The final quantity of each metal was normalised using dry weight biomass of the cell pellet prior to nitric acid digestion, expressed as µg.g^-1^dry weight.

### RNA sequencing and bioinformatics

Cultures were prepared as described above for *RNA extraction, qRTPCR* to compare mid-log phase WT or Δ*copY* cells grown in THB + 0.25 mM Zn or in THB without added Zn. RNase-free DNase-treated RNA that passed Bioanalyzer 2100 (Agilent) analysis was used for RNA sequencing (RNA-seq) using the Illumina NextSeq 500 platform. We used a Bacterial Ribosomal RNA (rRNA) Depletion kit (Invitrogen) prior to library construction, and TruSeq library generation kits (Illumina, San Diego, California). Library construction consisted of random fragmentation of the RNA, and cDNA production using random primers. The ends of the cDNA were repaired and A-tailed, and adaptors were ligated for indexing (with up to 12 different barcodes per lane) during the sequencing runs. The cDNA libraries were quantitated using qPCR in a Roche LightCycler 480 with the Kapa Biosystems kit (Kapa Biosystems, Woburn, Massachusetts) prior to cluster generation. Clusters were generated to yield approximately 725K–825K clusters/mm^2^. Cluster density and quality was determined during the run after the first base addition parameters were assessed. We ran single-end 75–bp sequencing runs to align the cDNA sequences to the reference genome. For data preprocessing and bioinformatics, STAR (version 2.7.3a) was used (parameters used: --outReadsUnmapped Fastx --outSAMtype BAM SortedByCoordinate -- outSAMattributes All) to align the raw RNA sequencing fastq reads to the WT *S. agalactiae* 874391 reference genome [55]. HTSeq-count, version 0.11.1 (parameters used: -r pos -t exon -i gene_id -a 10 -s no -f bam), was used to estimate transcript abundances [57]. DESeq2 was then used to normalized and test for differential expression and regulation following their vignette. Genes that met certain criteria (i.e. fold change of > ±2.0, q value (false discovery rate, FDR of <0.05) were accepted as significantly altered [58]. Raw and processed data were deposited in Gene Expression Omnibus (accession no. GSE167895 for *S. agalactiae* 874391 Cu condition; GSE167894 (*S. agalactiae* 874391 control condition).

### Animals and Ethics statement

Virulence was tested using a mouse model of disseminated infection based on intravenous challenge with 10^7^ GBS (WT, Δ*copY* or Δ*covR*) as described elsewhere [59]. This study was carried out in accordance with the guidelines of the Australian National Health and Medical Research Council. The Griffith University Animal Ethics Committee reviewed and approved all experimental protocols for animal usage according to the guidelines of the National Health and Medical Research Council (approval: MSC/01/18/AEC).

### Transposon Directed Insertion Site Sequencing (TraDIS)

Generation and screening of the 874391:IS*S1* library was performed essentially as previously described [60], with some modifications. Briefly, the pGh9:IS*S1* plasmid (provided by A. Charbonneau *et al*.) was transformed into WT *S. agalactiae*, and successful transformants were selected by growth on THB agar supplemented with 0.5μg/mL Erythromycin (Em). A single colony was picked and grown in 10mL of THB with 0.5μg/mL Em at 28°C overnight. The overnight cultures were incubated at 40°C for 3h to facilitate random transposition of IS*S1* into the bacterial chromosome. Transposon mutants were selected by plating cultures onto THB agar supplemented with Em and growing overnight at 37°C. Pools of the transposon mutants were harvested with a sterile spreader and stored in THB supplemented with 25% glycerol at -80°C. The final library of approximately 470,000 mutants was generated by pooling two independent batches of mutants.

Exposure of the library used approximately 1.9 x 10^8^ bacteria inoculated into 100mL of THB (non-exposed Ctrl) or THB supplemented with 1mM Zn in THB. The cultures were grown for 12h at 37°C (shaking), and subsequently, 10mL of culture were removed and washed once with PBS. Genomic DNA was extracted from three cell pellets per condition (prepared as independent biological samples) using the DNeasy UltraClean Microbial Kit (Qiagen) according the manufacturer’s instructions, except that the cell pellets were incubated with 100 units of mutanolysin and 40mg of RNase A at 37°C for 90min. Genomic DNA was subjected to library preparation as previously described [60], with slight modifications. Briefly, the NEBNext dsDNA fragmentase (New England BioLabs) was used to generate DNA fragments in the range of 200-800bp. An in-house Y-adapter was generated by mixing and incubating adaptor primers 1 and 2 (100μM, Supplementary Table S3) for 2min at 95°C, and chilling the reaction to 20°C by incremental decreases in temperature by 0.1°C. The reaction was placed on ice for 5min, and ice cold ultra-pure water was added to dilute the reaction to 15μM. The Y-adaptor was ligated to the ends of the fragments using the NEBNext Ultra II DNA Library Prep Kit for Illumina (New England BioLabs) according to the manufacturer’s instructions. All adaptor ligated fragments were incubated with *Not*I.HF (New England BioLabs) for 2h at 37°C to deplete plasmid fragments. The digested fragments were PCR amplified as per the protocol outlined in the NEBNext Ultra II DNA Library Prep Kit using a specific IS*S1* primer and reverse indexing primer (Dataset S4). DNA quantification was undertaken using a QuBit dsDNA HS Assay Kit (Invitrogen) and purified using AMPure XP magnetic beads (Beckman Coulter). All libraries were pooled and submitted for sequencing on the MiSeq platform at the Australian Centre for Ecogenomics (University of Queensland, Australia).

The sequencing data generated from TraDIS libraries were analysed used the Bio-TraDIS scripts [61] on raw demultiplexed sequencing reads. Reads containing the transposon tag (CAGAAAACTTTGCAACAGAACC) were filtered and mapped to the genome of WT *S. agalactiae* 874391 using the bacteria_tradis script with the “--smalt_y 1” and “--smalt_r 0” parameters to ensure accuracy of insertion mapping. Subsequent analysis steps to determine log_2_ fold-change (log_2_FC), false discovery rate (FDR) and P value were carried out with the AlbaTraDIS script [62]. To identify genes in *S. agalactiae* 874391 required for resistance to Zn intoxication condition used, we used a stringent criteria of log_2_FC ≤ -2 or ≥ 2, FDR <0.001 and P value <0.05. The TraDIS reads are deposited in the Sequence Read Archive (SRA) under BioProject ID: PRJNA674399.

### Statistical methods

All statistical analyses used GraphPad Prism V8 and are defined in respective Figure Legends. Statistical significance was accepted at P values of ≤0.05.

## Acknowledgments

We gratefully acknowledge Andrew Waller and Amy Charbonneau, Animal Health Trust (Suffolk, UK) for providing pGh9-IS*S1*. We thank Michael Crowley and David Crossman of the Heflin Center for Genomic Science Core Laboratories, University of Alabama at Birmingham (Birmingham, AL) for RNA sequencing. We also thank Lahiru Katupitiya and Dean Gosling for excellent technical assistance. This work was supported by a Project Grant from the National Health and Medical Research Council (NHMRC) Australia (APP1146820 to GCU). The funders had no role in study design, data collection and analysis, decision to publish, or preparation of the manuscript. The authors have declared that no competing interests exist.

## Author Contributions

M. J. S., G. C. U. conceived and designed the research. M. J. S., K. G. K. G., G. C. U. constructed the *S. agalactiae* mutants, M. J. S. performed the transcriptomic analyses, ICP-OES, phenotypic assays, cell and animal experiments and microscopy. K. G. K. G. performed the TraDIS experiments. M. J. S., K. G. K. G., G. C. U. discussed the results and wrote the manuscript together. All authors reviewed and edited the manuscript.

